# Phenotype-specific information improves prediction of functional impact for noncoding variants

**DOI:** 10.1101/083642

**Authors:** Corneliu A. Bodea, Adele A. Mitchell, Alex Bloemendal, Aaron G. Day-Williams, Heiko Runz, Shamil R. Sunyaev

**Author notes:** These authors jointly supervised this work (Aaron G. Day-Williams, Heiko Runz, Shamil R. Sunyaev).

## Abstract

Functional characterization of the noncoding genome is essential for the biological understanding of gene regulation and disease. Here, we introduce the computational framework PINES (Phenotype-Informed Noncoding Element Scoring) which predicts the functional impact of noncoding variants by integrating epigenetic annotations in a phenotype-dependent manner. A unique feature of PINES is that analyses may be customized towards genomic annotations from cell types of the highest relevance given the phenotype of interest. We illustrate that PINES identifies functional noncoding variation more accurately than methods that do not use phenotype-weighted knowledge, while at the same time being flexible and easy to use via a dedicated web portal.

## Introduction

A growing body of evidence suggests that DNA variants outside of protein-coding regions of the genome (here termed noncoding variants) impact human phenotypes, including the risk for common diseases. Many signals identified by genome-wide association studies (GWAS) reside in the noncoding genome and are hypothesized to have regulatory functions. Likewise, some human Mendelian phenotypes such as hair, skin, and eye pigmentation are under the tight control of individual highly penetrant noncoding alleles [1, 2]. Thanks to whole genome sequencing (WGS), our ability to uncover novel noncoding alleles has increased substantially. However, studies of both common and rare phenotypes almost never resolve findings in noncoding regions to individual, causal SNPs [3]. Statistical prioritization of noncoding variants thus holds significant promise to help interpret and prioritize variants identified in GWAS fine-mapping efforts and identify genetic lesions underlying Mendelian diseases.

To better understand the architecture of the noncoding genome, several large-scale efforts, such as the Encyclopedia of DNA Elements [4] and the NIH Roadmap Epigenomics Mapping Consortium [5], have aimed to characterize the diverse landscape of histone modifications and DNA accessibility based on a wide range of assays across 127 cell types. Other efforts such as the FANTOM5 project [6] have identified enhancer elements across the genome, and computational tools such as TargetFinder [7] have been developed to link such enhancers to the relevant gene promoters. Databases of chromatin interactions such as 4DGenome [8] are also helpful in identifying potential regulatory elements. Yet, while most genomic regions are by now annotated with a plethora of epigenetic features, the challenge remains to draw meaningful conclusions from these annotations, especially since the data are highly dimensional and many epigenetic features are correlated.

Several approaches have recently been introduced to prioritize potentially functional noncoding variants and address the complexity of the annotation data in a principled manner. These include Eigen-PC [9], GWAVA [10], CADD [11], DANN [12], FATHMM-MKL [13], LINSIGHT [14], deltaSVM [15], DIVAN [16], GenoCanyon [17], ReMM [18], and hyperSMURF [19]. One drawback of models such as GWAVA and deltaSVM is that they require a training dataset of both functional and non-functional examples. However, high-quality and particularly experimentally validated training data sets for noncoding variants are still very scarce and incomplete, thus limiting the prediction accuracy. A second drawback is that many existing scoring methods globally merge annotations across many different cell types, which ignores the observation that regulatory elements can operate in a very cell type-specific manner [20]. In addition, most methods that allow for a cell type-specific analysis, such as deltaSVM, do so by restricting their annotation sources or training set to solely a cell type of interest. This approach has multiple downsides. First, potentially informative annotations from similar cell types are ignored, which can decrease statistical power. Second, if the correct cell type is unknown, then focusing the analysis on a predefined cell type may lead to results that are not informative of the underlying biology. Finally, a large computational burden is placed on users by requiring them to provide annotations and training data from cell types of interest in addition to running the scoring method.

We hypothesized that in order to prioritize noncoding variants of relevance to a specific disease or trait, a principled integration of prior genomic information around the phenotype should increase prediction reliability. Such prior information can consist of phenotype-relevant cell types, tissues, genes or pathways. We further hypothesized that predicting the functional relevance of noncoding variants could benefit from taking into account the vast number of variants that have not yet been annotated as having any function (background variants), and use these as a baseline to search for variants that deviate significantly from this background. While the absence of reported functional data does not necessarily imply that all background variants stem from non-functional sites, this collection of variants can still serve as a useful reference against which to compare individual variants of interest.

Here we introduce the Phenotype-Informed Noncoding Element Scoring (PINES) framework. PINES enables a phenotype-dependent scoring of noncoding variants that can be customized towards annotations considered as of highest relevance to a phenotype of interest. During the PINES scoring process, such annotations are assigned the highest weights, while annotations of lower relevance to a phenotype are down-weighted. Moreover, differentiating from most previous scoring tools, PINES uses an unsupervised approach to systematically assess the functional significance of noncoding SNVs and indels. In particular, no variant sets are used to train the PINES model in the unweighted and manually weighted settings. When no phenotype-relevant information is available, PINES scores individual variants solely by comparing them to the background level of annotations across the genome, while taking the correlation structure between annotations into account. When the user specifies individual annotations as phenotype-relevant, this information is directly reflected in the annotation weights. Additionally, PINES generates weights in a data-driven process when supplied with a set of phenotype relevant variants (e.g. GWAS lead SNPs). Under this semi-supervised setting, the specified variants are only used to aid in the selection of annotation weights: after an enrichment analysis is performed independently for each annotation (see “Choice of feature weights” section under Methods) no further use is made of the variants, and the variants are never processed through the full PINES framework or scored.

PINES is a purely epigenetic-based noncoding variant prioritization approach and does not incorporate GWAS summary statistics directly into its scoring procedure, as opposed to methods such as PAINTOR [21] and fGWAS [22] which are meant as GWAS fine-mapping tools. PINES can thus help interpret the function of noncoding variation for complex phenotypes that do not yet benefit from extensive GWAS information, or for Mendelian phenotypes, for instance to shortlist rare noncoding variants of putative large effect that could help explain etiologies of rare disease. This is performed via upweighting user-specified annotations. PINES can also signal the presence of regulatory elements that are not yet tied to particular phenotypes. This is performed via unweighted scoring. To the best of our knowledge, no alternative variant interpretation tool can integrate known functional information into the scoring procedure, and also perform well when no prior phenotypic information is available.

We demonstrate the accuracy and versatility of PINES by addressing frequent challenges across several fields, for instance, predicting functional variants at assumed promoter and enhancer regions, prioritizing noncoding variants at GWAS loci, or assessing the relevance of noncoding variation on Mendelian phenotypes. With its ease of use via a customizable web server (http://genetics.bwh.harvard.edu/pines/) and open source code, we hope PINES will become a valuable resource for the interpretation of the noncoding human genome.

## Results

### PINES integrates epigenetic information to score non-coding variants in phenotype-centered queries

Fig.1 presents an overview of the PINES framework. PINES integrates data from each of the 127 cell types analyzed by the NIH Roadmap Epigenomics Mapping Consortium and the Encyclopedia of DNA Elements (ENCODE) project, specifically information on histone modifications indicative of promoters or enhancers (H3K4me1, H3K4me3, H3K27ac, H3K9ac), DNase I hypersensitive sites, sequence constraint scores (GERP, SiPhy), and ChromHMM chromatin state segmentations [23]. These data are applied to compare individual or a set of user-defined SNPs relative to a genomic background. When no prior knowledge is provided or available, PINES can be run in the default (unweighted) mode where equal weights are assigned to all features to perform an undirected scoring of variants. When prior information is provided as a list of GWAS lead SNPs associated with the trait of interest, PINES searches for annotations that are enriched in the provided list in order to learn which annotations are most relevant for the phenotype under consideration. Alternatively, users can manually specify the most phenotypically relevant annotations to be up-weighted by the scoring procedure. This allows that, if available, prior knowledge can be taken into account during the subsequent scoring process (see Fig. 1 and Methods for details).

**Figure 1.**
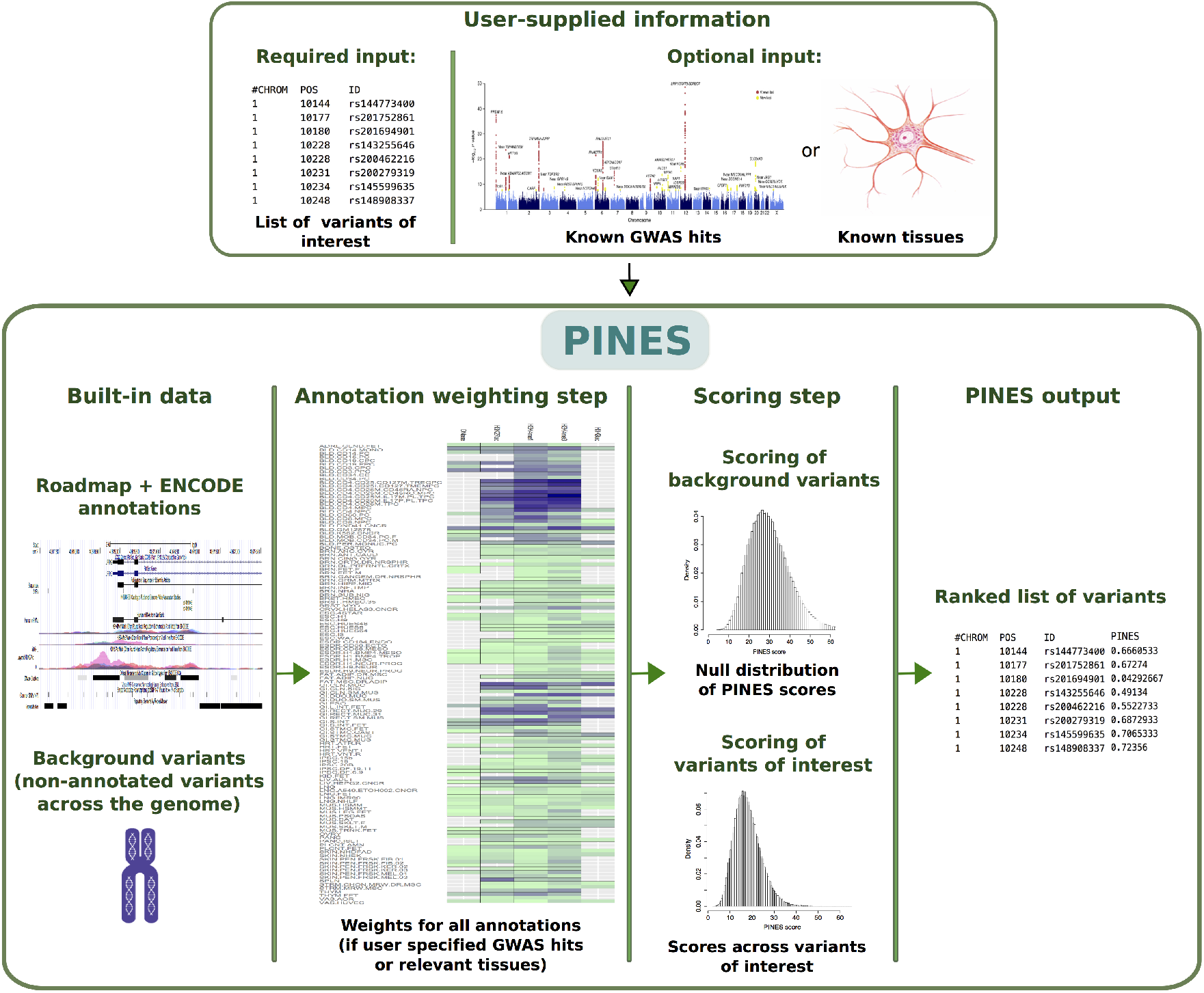
Overview of the PINES framework. PINES aims to systematically predict and rank the functional relevance of noncoding genomic variants. It can either work in a default (“unweighted”) mode and compare user-defined variants against the genomic background. Alternatively, users can customize searches towards annotations considered as of highest relevance to a phenotype of interest, for instance by providing a list of SNPs associated with a disease of interest through GWAS, or by highlighting disease-relevant tissues (‘weighted” PINES mode). Scores of genomic background variants serve as an empirical null distribution against which significance levels for each variant of interest are computed and scored in an output file.

The results of the ENCODE and NIH Epigenome Roadmap projects have produced a highdimensional, highly correlated representation of the genome that needs to be reduced and decorrelated in order to not over-fit or over-weight information when predicting functional variants. PINES addresses the high-dimensionality and high-correlation of the epigenetic annotations by developing an angle-based distance from the vector of the maximal possible annotation load in a de-correlated annotation space (see Methods). Prior genomic knowledge in the form of lead SNPs or pre-specified tissues and cell types is encoded as a weight matrix in PINES, where each annotation from each cell type receives an individual weight. When GWAS lead SNPs are provided, enrichment within GWAS peaks for each annotation is used to set weights (see section ‘Choice of feature weights” under Methods). As an example, the annotation weight matrices that PINES computes based on enrichment of epigenetic marks across inflammatory bowel disease, multiple sclerosis, celiac disease and blood lipids GWAS lead SNPs are shown in Supplemental Figs. S8 - S11.

A key conceptual advance of PINES (and a significant methodological advance over other methods) is its ability to conduct analyses in a user-defined manner, i.e., in a ‘weighted” mode. With the option to customize queries by a user, “weighted” PINES exceeds the accuracy of “unweighted” PINES as well as all other available tools tested for all datasets we have assessed. Feature weights represent a crucial component and add strongly to the power of PINES beyond other tools. The PINES framework can be used to identify functional noncoding variants in both Mendelian and complex disease settings.

### PINES prioritizes causal noncoding variants with high confidence

We first illustrate the potential of PINES to prioritize variants across noncoding genomic regions that have been directly implicated in Mendelian or complex traits. In order to test whether PINES correctly identifies and prioritizes known functional noncoding SNPs, we applied the algorithm to in silico score variants across 20kb regions around six noncoding alleles whose regulatory impact on a nearby gene have been confirmed experimentally: rs12821256 [24], rs6801957 [25], rs356168 [26], rs2473307 [27], rs12350739 [28], and rs227727 [29] (Fig. 2 and Supplemental Table S1). Such well-studied noncoding variants, although still rare in the current literature, provide an optimal testbed for scoring methods. Weighted PINES was used when genome-wide significant SNPs were identified by GWAS for the trait under consideration, and manual weighting was used for the pigmentation loci since, to our knowledge, no GWAS of pigmentation exists to date that has identified enough peaks for a well-powered enrichment analysis (see Methods). In all cases, PINES scores peak around the experimentally confirmed functional variants, and PINES assigns these causal variants the lowest p-values. Notably, comparative analyses using these regions demonstrate that PINES outcompetes GWAVA, CADD or Eigen-PC in correctly identifying causal noncoding variants (see Supplemental Figs. S1-S6). “Weighted PINES”, i.e., when PINES was run in a phenotype-weighted mode, differentiated causal variants from background variants even more clearly than when the software was run in the default (“unweighted”) mode, showcasing the additional benefits that come from including phenotype-relevant information into the scoring procedure.

**Figure 2.**
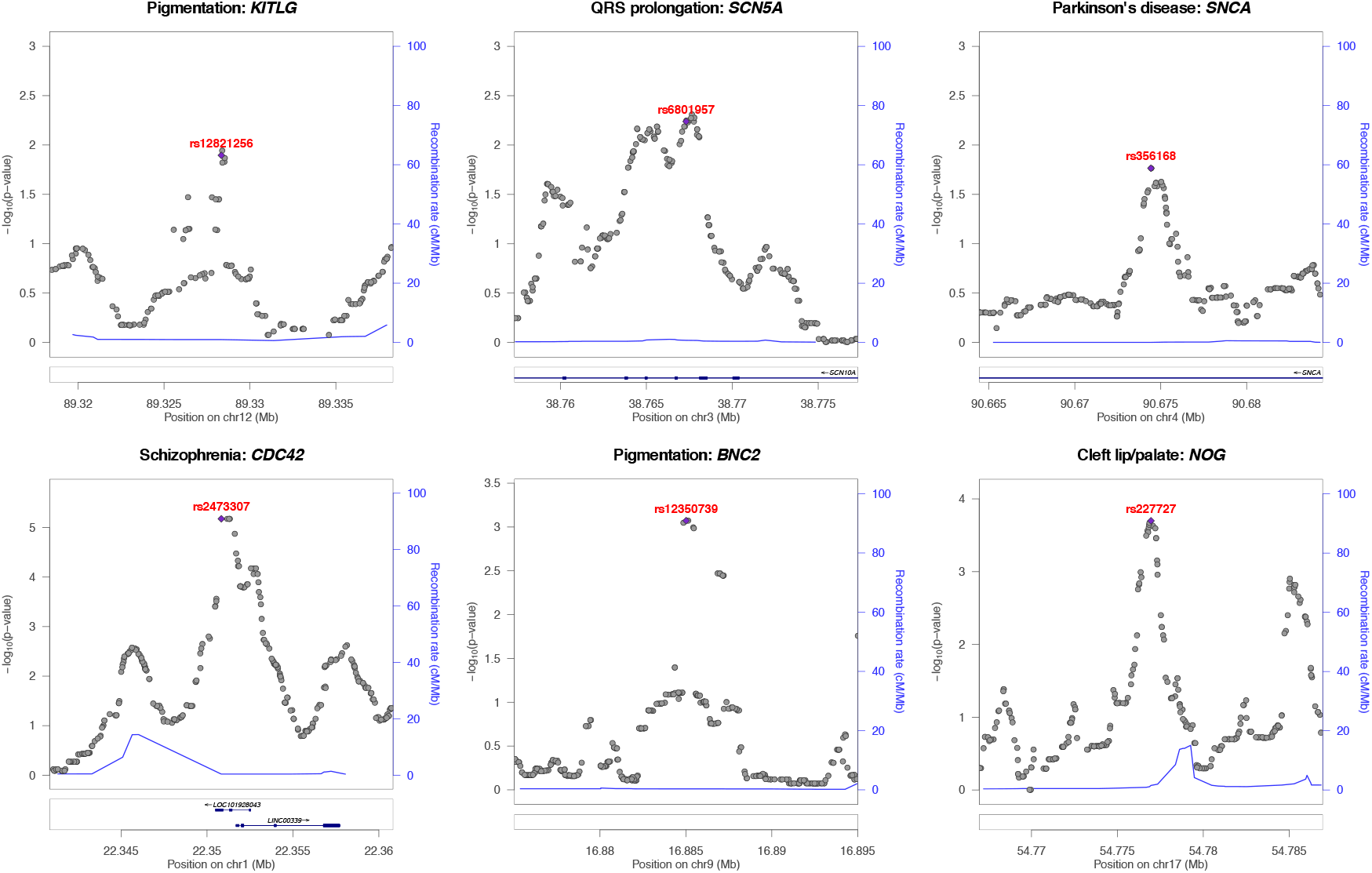
PINES prioritizes experimentally validated functional noncoding variants. We score all variants across 20kb regions surrounding functional noncoding variants (purple dots) and show that all of the variants validated experimentally as regulating expression of a nearby trait-associated gene are also assigned the highest PINES scores. Supplemental Figs. S1-S6 show that PINES outperforms existing methods on all loci.

### PINES detects signal at fine-mapped GWAS risk loci

We next aimed to evaluate how genomic information on the phenotype can improve the ability of PINES to predict causal noncoding variants at GWAS loci across a range of conditions. As a first example for this, we extracted 3,625 candidate causal noncoding variants (PICS probability ≥ 0.1) reported by a large statistical fine-mapping study spanning 21 autoimmune diseases as well as some non-immune diseases [30]. This study performed fine-mapping by using densely-mapped genotyping data and the observed pattern of association at the LD locus to estimate each SNP’s probability of being a causal variant. For the fine-mapped variants associated with immune phenotypes we computed weighted PINES scores, where all annotations corresponding to immune cells (primary T and B cells) were equally up-weighted (see Methods). In this use case, PINES weights are thus specified manually and not learned from GWAS data. We compared the collective PINES signal on these variants with the results obtained on an additional set of 30,000 background variants that we randomly selected across the genome. Q-Q plots detailing the results highlight a well-calibrated PINES null distribution and strong signal on the fine mapped variant set (Supplemental Fig. S7). Comparison of the weighted PINES results with outcomes from GWAVA, Eigen-PC, CADD, LINSIGHT, DANN, DIVAN, GenoCanyon, and FATHMM-MLK shows that PINES delivers the best predictive performance (Fig. 3 first panel). Also to note is that only 37% of the candidate causal noncoding variants identified by PICS also obtain a weighted PINES p-value ≤ 0.05. This highlights the ability of PINES to prioritize results from fine-mappixng studies to further restrict the search space for causal variants and enable focused experimental follow-up.

**Figure 3.**
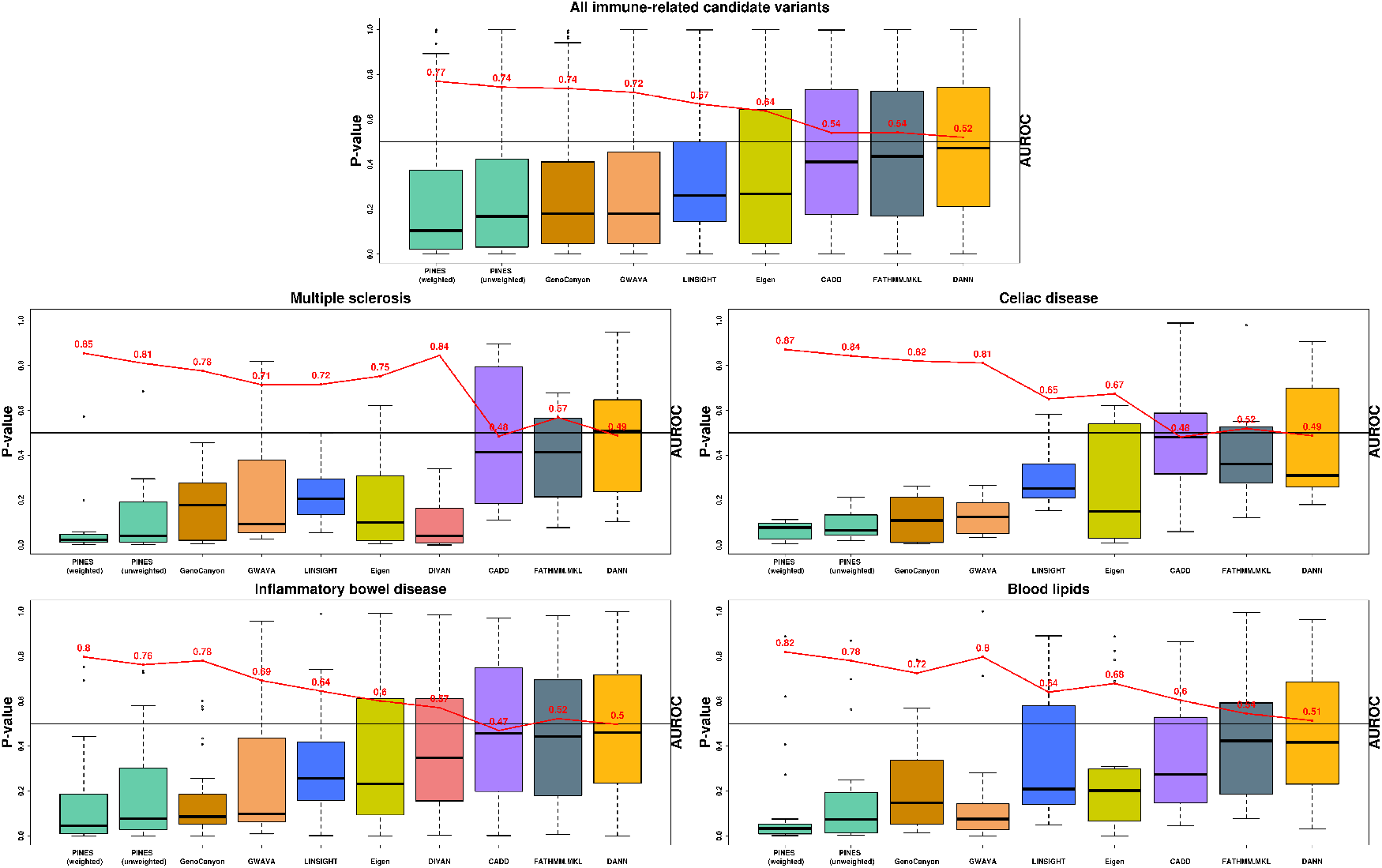
PINES improves statistical power to detect fine mapped variants across common neurologic, immune, and metabolic traits and diseases. AUROC values (red) were computed by selecting 30,000 background variants as negative examples, and the fine mapped variants relevant to each disease as positive examples. Weighted PINES consistently achieves better classification accuracy than the other methods, based on its inclusion of weights encoding prior disease knowledge (relevant cell types or GWAS lead SNPs).

Next, we assessed the performance of PINES to nominate causal alleles from loci associated with individual complex traits and diseases, including non-immune phenotypes. We extracted high-confidence (posterior probability ≥ 0.25) fine mapped candidate causal noncoding variants from [30] related to multiple sclerosis (12 variants), celiac disease (7 variants), inflammatory bowel disease (31 variants), and blood lipid levels (22 variants). We determined weighted and unweighted PINES scores for each of these variants, and compared the outcomes of PINES to those of GWAVA, Eigen-PC, CADD, LINSIGHT, DANN, DIVAN, GenoCanyon, and FATHMM-MLK. Phenotype-based annotation weights were automatically assigned based on GWAS data ( [31–36]). PINES consistently delivered the highest AUROC values when run in the phenotype-weighted mode (Fig. 3). Upweighting of phenotype-relevant annotations also enabled weighted PINES to outcompete unweighted PINES results on all individual phenotypes.

Of note is that we notice a drastic improvement in classifier accuracy when PINES computes annotation weights based on GWAS data in the MS, Celiac disease, IBD, and blood lipids analyses. The panel highlighting all immune-related candidate variants groups together variants across many immune disorders, and thus relies on a uniform manual up-weighting of all cell types involved in immunity (primary T and B cells). Classification accuracy is increased in this setting as well compared to an unweighted analysis, but an enrichment-based weighting of features is preferred.

Since we expect to see overlap between the GWAS lead SNPs (used to construct the annotation weights) and the high-confidence (posterior probability ≥ 0.25) PICS variants that resulted from fine-mapping of these GWAS loci, we formally evaluated the effect of this overlap on our comparisons. To note is that GWAS lead SNPs are only used to extract a high-level representation of the relevant cell types for weighting, but are not carried forward into the scoring method. Supplemental Figs. S8-S11 depict all the information that is extracted from these variants and used in the subsequent weighted scoring step. These figures show that PINES upweights biologically relevant cell types for each disease (IBD: immune and GI, MS: immune and thymus, celiac disease: immune and GI, blood lipids: liver and adipocytes). In addition, these GWAS lead SNPs are only used in an aggregated manner by evaluating the overlap of the entire set of variants with epigenetic annotations (a detailed description of the weighting algorithm is presented in the “Choice of feature weights” section under Methods). Thus, no analysis is performed to learn features of individual GWAS lead SNPs.

We used two different approaches to test whether the presence of variants used both as GWAS lead SNPs (to compute annotation weights) and scored PICS SNPs introduces any bias in our comparisons. First, within the set of variants that achieved high PICS posterior probabilities (posterior probability ≥ 0.25) we evaluated whether also being a GWAS lead SNP resulted in more significant weighted PINES scores. We performed a Wilcoxon rank-sum test comparing the weighted PINES scores of PICS variants that are also GWAS lead SNPs versus those that are not GWAS lead SNPs, and observed no significant difference in the distribution of scores between these two categories (MS: p-value = 0.7849, IBD: p-value = 0.7678, blood lipids: p-value = 0.6311, celiac disease: p-value = 0.7302). Second, we re-ran all four disease-centered comparisons in Figure 3, but excluded all PICS variants that were also used in the computation of cell type weights. This way we only use variants that PINES has never seen in the cell type analysis step. The performance of PINES remains unchanged in comparison to the other methods: weighted PINES remains the best performing method based on the resulting AUROC values.

### Unweighted PINES scores detect signal at expression-modulating variants identified in a multiplexed reporter assay, and for variants residing in FANTOM5 CAGE-defined enhancers

To compare the performance of PINES, Eigen-PC, CADD, GWAVA, DANN, LINSIGHT, Geno-Canyon, and FATHMM-MLK to detect functional noncoding variants en masse when no phenotype information is available to guide the scoring, we used 230 variants that were found to directly affect gene expression in a massively parallel reporter assay (MPRA) [37]. For the selection of variants we used an FDR cutoff (Benjamini-Hochberg) of 1%. MPRA is an extension of the traditional reporter gene setup, whereby the use of unique barcodes in the 3’ UTR of the reporter differentiate expression of individual oligos and thus allow for testing of many different sequences simultaneously. Since most variants identified through this approach have not yet been linked to individual phenotypes, we performed an unweighted PINES analysis. PINES outscored all other methods except for GWAVA and GenoCanyon, which delivered similar results (Fig. 4).

**Figure 4.**
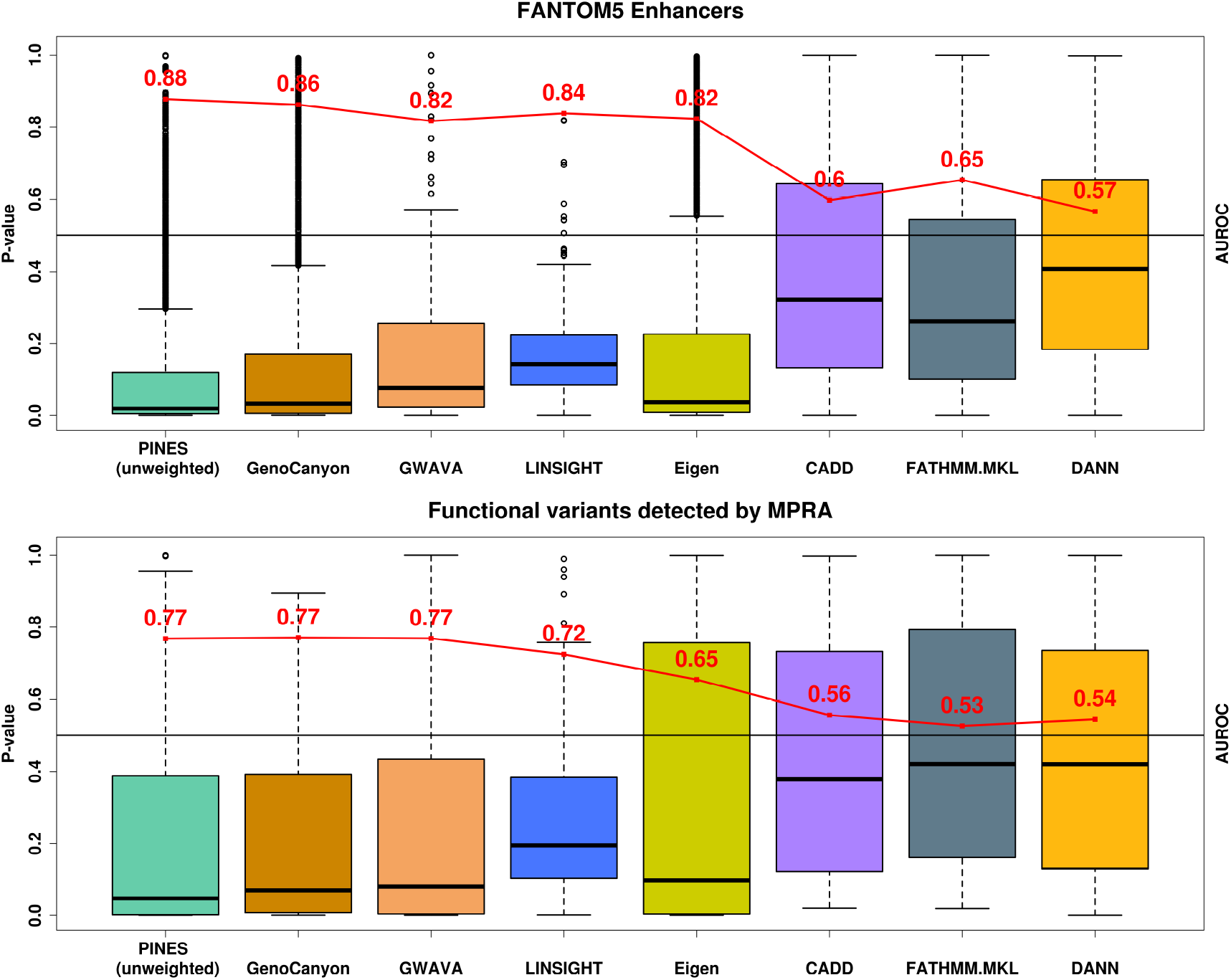
Unweighted PINES scores improve the prioritization of noncoding variants. (A) PINES improves the prioritization of variants residing in experimentally validated enhancer regions. The AUROC values (red) were computed by selecting 20,000 background variants as negative examples, and the variants residing in enhancer loci as positive examples. Based on AUROC values, the unweighted PINES approach performs at least as well as GWAVA, Eigen-PC, CADD, DANN, LINSIGHT, GenoCanyon, and FATHMM-MKL in its ability to pinpoint enhancer variants. (B) PINES delivers improved statistical power to identify functional noncoding variants detected by a massively parallel reporter assay. The AUROC values (red) were computed by selecting 20,000 background variants as negative examples, and the reported functional variants as positive examples. PINES achieves better classification accuracy than the other methods, outperforming GWAVA, Eigen-PC, CADD, DANN, LINSIGHT, GenoCanyon, and FATHMM-MKL in its ability to detect the functional variants.

We next compared the power of PINES, Eigen-PC, CADD, GWAVA, DANN, LINSIGHT, GenoCanyon, and FATHMM-MKL to correctly prioritize 9,000 variants residing within enhancers that have been identified by the FANTOM5 project through cap analysis of gene expression (CAGE) [6]. Since no phenotype has been recorded for the variants residing in FANTOM5 enhancers or for the MPRA hits, we only performed an unweighted PINES analysis. Thus no tissue-specific information was included in the scoring process. We used all CAGE-based human enhancers reported by FANTOM5 (phases 1 and 2 combined). When defining our set of background variants, we found 9,000 variants that overlapped with these enhancer regions (see section “Background variants and the null distribution” under Methods). We excluded these variants from the background set, and instead scored them via PINES to investigate whether any enrichment in epigenetic signal can be detected relative to the genome-wide background. PINES correctly assigned noncoding variants residing in these regions the highest relevance scores relative to genomic background variation (Fig. 4).

These analyses highlight that, even in settings where no phenotype-relevant information is available, none of the methods that we tested were able to perform better than PINES. Reasons for the good performance of PINES are the unsupervised nature of the framework that generates unweighted scores (no training set with an explicit focus on disease-associated variants is used), as well as the use of a de-correlated search space and angle-based outlier detection approach (see Methods).

### PINES identifies variants with highly cell type-specific annotation profiles

A comparison of weighted and unweighted PINES scores on the same set of variants can pinpoint noncoding variants that exhibit epigenetic annotations in a cell type-specific manner. As an example, we show that such an analysis can be useful in identifying variants with GI-specific annotation profiles in IBD.

We simulated 5000 background variants and 100 cell type-specific variants across 100 binary annotations, and randomly selected 10 annotations for up-weighting. We simulate binary annotations since they depict whether a variant overlaps or does not overlap with a given epigenetic mark. For the cell type-specific variants, the up-weighted annotations are drawn from a Bernoulli distribution with p=0.9 and the remaining annotations are drawn from a Bernoulli distribution with p = 0.1. For the background variants, all annotations are drawn from a Bernoulli distribution with p = 0.3. The resulting cell type-specific variants have low unweighted scores and high weighted scores (Fig. 5A). As a quantitative measure of location in this score space, we compute the angle formed by each variant to the main diagonal. Cell-type specific variants lead to significantly larger angles than background variants (Fig. 5B), and can thus be identified. Applying this approach to lead SNPs from the inflammatory bowel disease GWAS [34] we can detect variants with, for example, GI-specific annotation profiles (Fig. 5C).

**Figure 5.**
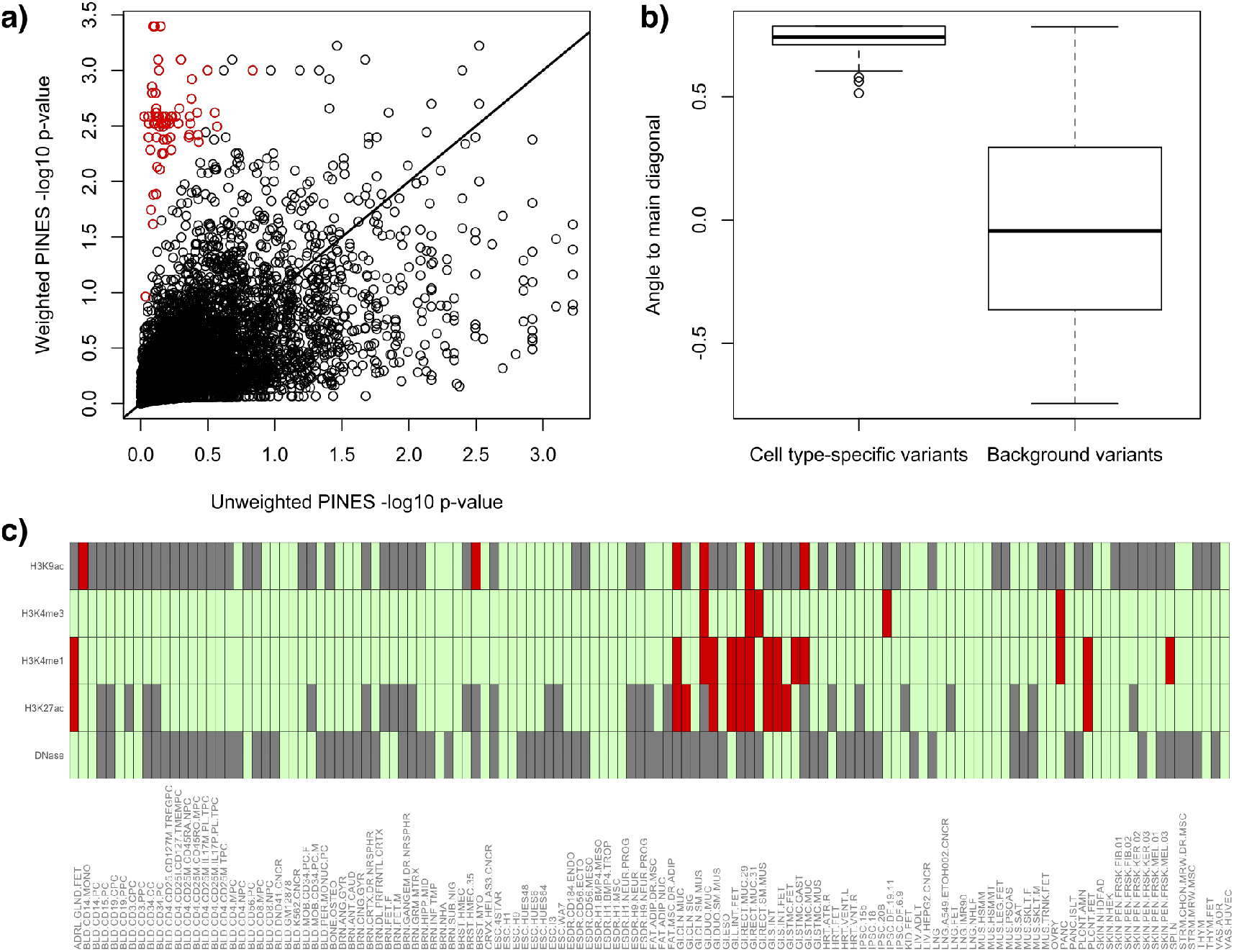
A comparison of weighted and unweighted PINES scores reveals cell type-specific variants. We simulate 5000 background variants (black circles) and 100 cell type-specific variants (red circles) and compute both weighted and unweighted PINES scores. Weighting is based on the annotations that are representative for the cell type-specific variants. The red dots are easily distinguishable from the background variants based on their location in the PINES score space (panel A) as well as the angles they form to the main diagonal (panel B). Applying this approach to variants from the inflammatory bowel disease GWAS in [34] we can detect noncoding variants with putative GI-specific activity. As an example, panel C depicts the annotations present at rs6017342 (red: annotation present, green: annotation absent, gray: missing data), a variant rich in GI-specific annotations that has been implicated by fine mapping of IBD GWAS loci [38].

### PINES points to novel causal variants for Parkinson’s disease and IBD through epigenetic prioritization at GWAS loci

We next tested whether phenotype-weighted PINES can be applied to predict novel functional noncoding SNPs from GWAS loci. For this, we applied PINES to all GWAS loci associated in recent meta-analyses at genome-wide significance with Parkinson’s disease [39] and IBD [34]. We concentrated on those loci where all SNPs in LD of *R*^2^ ≥ 0.4 to the GWAS lead SNP were either intronic or intergenic. This resulted in 12 Parkinson’s disease loci and 19 IBD loci. It is likely that many more Parkinson’s disease and IBD loci are driven by noncoding mechanisms, however the lack of coding variants in the LD blocks surrounding our selected loci increases our confidence in a set of underlying causal noncoding variants. For both Parkinson’s disease and IBD we used PINES to determine enrichment-based phenotype-specific weights from the complete set of significantly disease-associated GWAS SNPs. We then ran weighted PINES to prioritize the most likely functional SNP across the selected Parkinson’s disease and IBD loci. With this approach, PINES significantly distinguished 16 noncoding alleles from the background, implicating them based on epigenetic grounds as likely causal for conferring risk for Parkinson’s disease (rs10878226, rs3756063, rs2301134, rs36121867, rs1954874, rs9275373, rs117896735) and IBD (rs35493230, rs2187892, rs4672507, rs4845604, rs2019262, rs10489630, rs12622128, rs55776317, rs7685642)(Fig. 6 and Supplemental Tables S2-S3).

**Figure 6.**
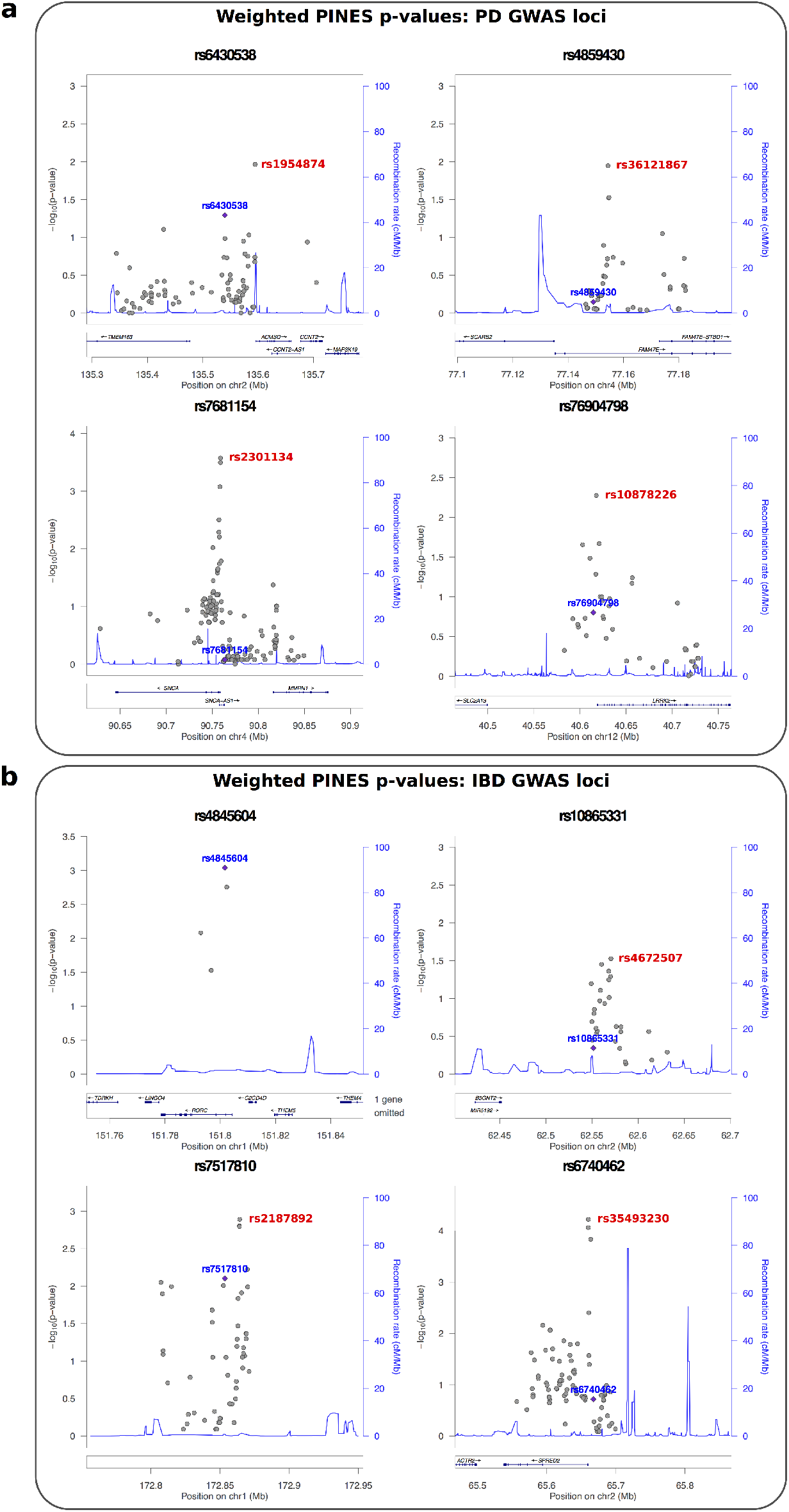
PINES predicts novel noncoding pathogenic variants through epigenetic prioritization of variants in Parkinson’s disease and IBD GWAS loci. Loci were extracted from [39] and [34]. For each lead SNP, all variants with *LD* ≥ 0.4 were selected, and loci were discarded if this list overlapped any coding regions or 3’ or 5’ UTRs of coding genes. All variants in LD to the lead SNP were scored via weighted PINES. The GWAS lead SNP is marked blue, and the variant predicted as likely causal through PINES prioritization is marked red. For the rs4845604 locus the GWAS and PINES lead SNP overlaps.

## Discussion

Human genetics is ever-increasingly pointing to an important role of noncoding variants in complex and Mendelian disease. However, which of these variants are important and in what context the consequences of their effect are exerted remain elusive. Hence, there is a strong need to shift the focus of the field from identifying novel signals towards better understanding what they actually mean. Yet, the notion of biological function of a variant is diffuse. Most genetically mediated common, complex traits are an interplay between a few but limited number of tissues or cell-types. Some of them are pleiotropic and affect several systems, but very few represent truly systemic disorders or traits that impact every cell in the body. An example is IBD, whose etiology points to both immune and gut epithelial mechanisms, and GWAS of IBD highlights both axes. This suggests that, from the genetic perspective, the notion of function only makes sense in the specific phenotypic context defined by cell type, developmental stage, and stimulus response. This is especially true for regulatory variants involved in transcriptional control. A number of recent studies showed that genetic association signals are enriched in putative regulatory elements, and that this enrichment is highly cell type-specific [30]. However, many experimental and almost all computational approaches to probe the functional effects of allelic variants are agnostic about the context of the phenotypic presentation. PINES has the unique ability not only to use prior genomic knowledge on the phenotype to customize the scoring procedure, but it can also identify the relevant cell types based on previous GWAS data, and can point out variants that exhibit annotation profiles that are specific to a cell type of interest.

Functional genomics is now actively embracing the multitude of contexts, starting from cell type variability in epigenetic annotations [5]. PINES leverages this annotation richness and attempts to predict the actual functional relevance in the most appropriate context rather than in the abstract framework of ubiquitous functional relevance. We note that simply restricting the analysis to the most relevant cell type is not the optimal approach. From a purely statistical perspective, noisy correlated data provide information and should not be completely neglected. More importantly, from the biological perspective, many alleles are pleiotropic and many phenotypes are influenced by different biological processes in different organs, tissues and cells. For example, risk of myocardial infarction is partly influenced by blood lipids, but many genetic contributions are unrelated to blood lipid levels and are likely mediated by the vascular effects. All autoimmune diseases are influenced by the adaptive immune system, but individual conditions are limited to specific organs. PINES can learn this complexity based on the given input information, and through its customizable weighting of annotations PINES can compute scores that take this context into account. Additionally, many cell types that are relevant to a phenotype are currently not represented in the ENCODE and Roadmap datasets. The ability of PINES to leverage information from related cell types and tissues enables the analysis of noncoding variants even for such phenotypes. Finally, the noncoding genome has been consistently linked to human phenotypes through our knowledge of conservation, GWAS peak localization, and eQTLs, yet so far only a few noncoding loci have been experimentally validated. This lack of unambiguously-defined functional noncoding loci makes the unsupervised approach used by PINES very versatile. By performing noncoding variant prioritization based solely on epigenetic annotations, PINES can aid in prioritizing noncoding variants underlying complex phenotypes that do not yet benefit from extensive and well-powered GWAS information, as well as prioritize lists of rare noncoding variants stemming from family studies of Mendelian phenotypes.

For users less experienced in working with the open-access PINES source code, PINES can be easily queried through a web server at http://genetics.bwh.harvard.edu/pines/ in a similar manner to our previous variant interpretation tool PolyPhen [40], which had been tailored to assess the consequences of coding variation. The web portal allows for scoring of noncoding SNVs based on user-defined weighting schemes, making PINES immediately applicable across a wide range of phenotypes. In addition, since the web server performs all data processing, users can query PINES with minimal computational overhead. PINES allows for the addition of epigenetic annotations as they become available without requiring significant changes to the underlying statistical model or software implementation. Due to this ease of upgrading the underlying annotation database, we aim PINES to become an always-up-to-date resource for the scientific community.

In conclusion, PINES’ ability to take advantage of a wide range of prior functional genomic information allows it to improve on the predictive power of other methods, and to provide an enhanced prioritization of phenotype-relevant variants. PINES avoids biases stemming from inaccurate labeling of training datasets, and benefits from increased power when prior information is available to direct analyses towards relevant annotations. There is a great need for such methods since systematic identification of regulatory activity specific to a subgroup of cell types or tissues can greatly increase our understanding of the still under-explored noncoding part of the human genome, as well as disease mechanisms linked to it.

## Methods

### Annotation sources

PINES uses a wide range of annotations as part of the scoring algorithm. Open chromatin and histone modifications for 127 cell types and tissues were obtained from ENCODE and Roadmap Epigenomics ChIP-seq and DNase-seq peak sets. In order to capture combinatorial interactions between different chromatin marks in their spatial context, we used ChromHMM chromatin state segmentations from Roadmap Epigenomics computed via the standard 15-state HMM model. Chromatin interaction data from a variety of assays (3C, 4C-Seq, 5C, Hi-C, ChIA-PET, Capture-C) were obtained from the 4DGenome database [8]. Additional DNaseI regions inferred via HMM from ENCODE and Roadmap Epigenomics data were obtained from the Reg2Map database. Conservation was evaluated via GERP [41] and SiPhy [42].

Noncoding background variants were selected randomly across the genome, and we used the ClinVar database [43] and GWAS Catalog [44] to ensure that no overlap exists with known functional loci (see section “Background variants and the null distribution” for a detailed description of the background variant set). For the analysis in Fig. 3 we used GWAS studies of IBD [33, 34], celiac disease [32], blood lipid levels [35, 36], and multiple sclerosis [31] to determine enrichment-based weights for weighted PINES. We then used fine mapped variants on the corresponding phenotypes from [29] as our test set. All immune-related fine mapped variants in [29] with posterior probability ≥ 0.1 were used to generate the first panel in Fig. 3. A list of CAGE-defined FANTOM5 enhancers [6] was used to create Fig. 4. The analysis of all noncoding Parkinson’s disease and IBD loci (Fig. 6) was based on regions identified in [39] and [34].

### Working with a high-dimensional correlated annotation space

Individual variants are assigned a score of 0 or 1 for each of the annotations referenced above. In particular, each variant is characterized by a vector of length 639 composed as follows:

- Presence or absence of H3K4me1, H3K4me3, H3K27ac, H3K9ac, and DNase annotations for each of the 127 epigenomes (635 values).
- Presence or absence of a conserved region as predicted by GERP and SiPhy (2 values).
- Presence or absence of a DHS region as predicted by the ChromHMM 15 state model trained on all epigenomes (1 value). This annotation represents ChrommHMM predictions that aggregate annotation across all cell types, so only one value is provided. However, cell-type specific DHS data is also included (see the first bullet point).
- Presence or absence of a region involved in chromatin interactions with other regions as reported in the 4DGenome database (1 value). Due to the substantial variability between the number of chromatin interactions reported for different cell types in the 4DGenome database, we aggregate the available chromatin interaction data across all cell types and treat this as one non-tissue-specific annotation.

In our annotation dataset, each variant is thus characterized by a vector of 635 cell type-specific scores and 4 cell type-independent scores. The joint distribution of this vector is difficult to ascertain explicitly due to its complex correlation structure; there are few outlier detection techniques that are robust to correlated data. One such approach is the one-class support vector machine (SVM) [45–47], which fits a hyperplane or hypersphere to the data in an attempt to isolate outlying points. One-class SVMs however suffer from a few disadvantages, such as difficulty in choosing tuning parameters, and the inability to add user-specified feature weights.

The alternative approach used in PINES is based on angular distances in a de-correlated annotation space. Let **X** be the 639-dimensional matrix of annotations with covariance matrix **Σ** and mean vector *μ*, and let **W** be a diagonal matrix of annotation weights (which in an unweighted analysis is the identity matrix). Since **Σ** is a noisy estimate of the true correlation structure of **X**, we perform the spectral decomposition 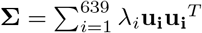, and compute a low-rank approximation of the estimated covariance matrix based on the first 30 eigenvectors (chosen by visual inspection of the scree plot): 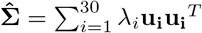. To account for potential differences in the annotation architecture of intronic and intergenic variants, we compute the background covariance matrix separately for these two variant classes. We noticed similar correlation structures, with the top 30 eigenvectors accounting for 53% of the variance for intronic variants, and 49% of the variance for intergenic variants. The matrix 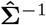 is obtained via the Moore-Penrose pseudo-inverse and is used to project annotation vectors corresponding to individual variants into a decorrelated annotation space via a Cholesky transformation: 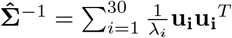. If x is a vector of annotations, then the length of the vector projected into the decorrelated space is 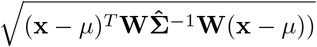, which corresponds to the reweighed Mahalanobis distance to the mean vector *μ* [48].

In summary, this formula allows us to project annotation vectors of individual variants into a decorrelated annotation space, and compute their distance to the mean annotation vector in this space. We also account for noise in the annotation covariance matrix via a low-rank approximation, account for differences between intronic and intergenic annotation profiles by considering these two types of variants separately, and enable the inclusion of weights into the scoring procedure. At this point we could thus use the actual value of 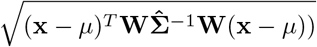 as a functionality score. Besides avoiding a double-count of correlated annotations, this score is also sensitive to variants whose annotation vectors break the overall correlation patterns. For example, if two annotations show a strong positive correlation overall, yet a variant only overlaps one of them, then this will be reflected in the score. While detecting deviations from established correlation patterns is an important feature, it can also lead to suboptimal results for variants that overlap many epigenetic annotations. To illustrate this effect, we generated a random correlation matrix for 100 annotations (Supplemental Fig. S16 panel A), and simulated 20,000 binary vectors (representing the individual variants and their overlap with each annotation) from these correlated annotations. Based on this simulated dataset, Supplemental Fig. S16 panel B shows a reduction in significance levels once variants approach overlap with all 100 annotations. This behavior is driven by the higher expectation (based on the positive correlations) that a variant will overlap many annotations or few annotations, rather than part of the annotations. In our scoring procedure we are however interested in also highlighting variants with high annotation load, so we introduce an angle-based scoring procedure in this de-correlated annotation space.

The cosine of the angle between the projections of two vectors x and y into the decorrelated annotation space is given by

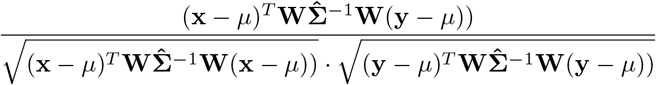

which corresponds to the correlation between the projections of x and y. In PINES we are specifically interested in the angle between the projection of a variant of interest and the projection of the all-1 annotation vector 1. This vector is significant since it provides the direction of a point with maximal annotation load. Under the assumption that biochemical marks such as histone modifications are indicative of variant functionality, the presence of multiple epigenetic marks indicative of regulatory regions that are supported across multiple cell types provides the greatest level of functional evidence for the all-1 annotation vector. Reference [49] presents an outlier detection method that addresses the curse of dimensionality by using an angle-based distance measure in addition to traditional Euclidean distance. We apply this principle in PINES, by scaling the angle between the projection of a variant of interest and the projection of the all-1 annotation vector by the length of the projected x vector. This results in the following PINES score:

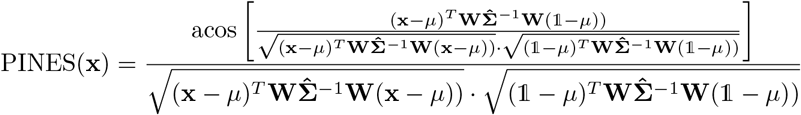

Smaller scores indicate greater evidence for functionality. Finally, to determine the significance level of a given score we compute an empirical p-value based on a large set of background variants. These background variants represent 150,000 common variant sites (since rare variant sites can potentially harbor more penetrant mutations) selected randomly across the genome and that are not represented in ClinVar or the GWAS Catalogue.

### Background variants and the null distribution

PINES makes use of a set of background SNPs against which to compare the score for new variants. The set of background variants was selected according to the following steps:

1. Extract a random subset of 200,000 rs IDs from dbSNP build 137. No additional criteria were used in the randomization process at this stage.
2. Annotate the resulting variant set with ClinVar data, and remove all “Pathogenic”, “Likely pathogenic”, and “Uncertain significance” variants.
3. From the resulting variant set, remove all variants that are present in the GWAS Catalog, or overlap FANTOM5 CAGE-based human enhancer regions.
4. Annotate the remaining variants with 1000 Genomes continental allele frequencies (AFR, AMR, ASN, EUR), and remove all variants with frequency ≤ 5% for any of the continental ancestries.

This stepwise filtering results in a background set of 155,000 variants. The median distance between two consecutive background SNPs is 5.7kb, which indicates a good coverage of the entire genome. We observe only 4,387 background variants (about 2.8% of the background set) that are in LD (*r*^2^ ≥ 0.8) with any GWAS Catalog variant. Our background set is also representative of a wide range of distances to the nearest transcription start site (TSS):

1. first 25% of variants: 0 kb - 25kb from the nearest TSS
2. second 25% of variants: 25kb - 87kb from the nearest TSS
3. third 25% of variants: 87kb - 256kb from the nearest TSS
4. fourth 25% of variants: more than 256kb from the nearest TSS

Since regions close to gene transcription start sites are enriched in epigenetic marks, our selection of background variants ensures a good match when scoring variants is such regions. PINES also automatically annotates all variants with their distance to the nearest TSS. This annotation is not used by default in PINES, but the user can enable it via the source code.

The number of background variants corresponding to each chromosome correlates well with chromosome length (Supplemental Fig. S12). The background variants also display a good coverage of the autosomal genome (Supplemental Fig. S13), with annotations across sex chromosomes not yet included in the PINES model.

Based on this background, PINES reports a one-sided p-value for each input variant, but the scores reported by CADD, GWAVA, and Eigen-PC do not have an absolute unit of meaning and are thus not directly comparable. To enable this comparison we use the collection of background variants to determine a null distribution for each scoring method, and transform the raw CADD, GWAVA, and Eigen-PC scores into empirical one-sided p-values based on their respective null distribution. This approach is similar to the one used to compute scaled CADD scores by transforming raw into rank-based scores [11]. Having performed this normalization, we can compare the results of PINES, CADD, GWAVA, and Eigen-PC directly.

### Choice of feature weights

When GWAS peaks are available, the weights used by PINES to differentiate between the different cell type-specific annotations are automatically computed based on the enrichment of each annotation across the GWAS loci. Enrichments are computed by performing 1000 draws of variant sets that are matched by allele frequency to the user-specified GWAS lead SNPs. We then compare the total number of GWAS variants overlapping each annotation with the number of overlaps in the matched sets to compute an empirical enrichment p-value. Enrichment within GWAS peaks for each annotation is used to set weights for the Parkinson’s disease, QRS prolongation, schizophrenia, and cleft lip and cleft palate variants presented in Fig. 2. Such enrichment-based approaches are frequently used in predicting cell types contributing to specific phenotypes when GWAS or fine mapping data is available [30], with highly enriched annotations indicating potentially relevant cell types and disease mechanisms. In particular, we used the corresponding -log10(enrichment p-value) as weight for every annotation, although different functional forms are possible. Supplemental Fig. S8 highlights the weights placed by PINES on cell-type specific annotations when IBD GWAS lead SNPs ( [34]) are provided as input by the user. To note is the high weight placed on immune and GI cells, two distinct cell groups that are known to be involved in IBD pathology. Supplemental Fig. S9 shows the weights placed by PINES on cell-type specific annotations when MS GWAS lead SNPs are provided as input by the user. To note is the high weight placed on immune cells, which is expected given the autoimmune etiology of MS. PINES also upweights annotations related to the thymus and fetal thymus, which have been connected to MS pathology as a source of autoreactive T-cells. Similarly, Supplemental Figs. S10 and S11 plot the weights placed by PINES on cell-type specific annotations when lead SNPs from GWAS of celiac disease and blood lipids are provided as input by the user.

Weighting of features can also be performed manually, as was the case for the pigmentation variants presented in Fig. 2, the first panel in Fig. 3, and the simulation in Supplemental Fig. S14, by setting the weight for biologically relevant annotations to a user-specified constant. Unweighted PINES scores are computed by setting the weight matrix W to the identity matrix (all weights = 1). The problem of manually up-weighting a subset of annotations is equivalent to specifying a constant to serve as weight for the annotations of interest. If this constant is set to 1, then we are in the unweighted case. If the constant is greater than 1, then more weight is placed on the corresponding annotations.

To determine an appropriate value for the weighting constant, it is important to note that increasing the value beyond a certain threshold results in two outcomes: the increase will stop adding power to detect variants overlapping the annotations of interest; and the increase will significantly reduce the power to detect variants overlapping other annotations. An appropriate weighting constant will provide sufficient power to detect variants overlapping annotations of interest while still allowing PINES to detect signal even if it does not stem from the upweighted annotations. While the choice of weighting constant will thus depend on the selected annotations, we have empirically observed via simulations that a weighting constant of 4 generally achieves a good balance of power given the correlations present in the ENCODE and Roadmap data.

Supplemental Fig. S15 formalizes this argument by highlighting the dynamics of weighted PINES scores in simulated data as the weighting constant increases. For this analysis we generate ten sets of correlated annotations. Each set contains ten binary annotations (0/1), and no correlation exists between annotations of different sets (panel a). We then select the first block of annotations as biologically relevant, and assign the same weighting constant to all ten annotations within the block. This simulates the presence of phenotypically-relevant annotations that the user selects to upweight. We compute the PINES significance level for two variants as the weighting constant increases from 0 to 20 in 0.5 increments: variant 1 overlaps all ten upweighted annotations and none of the remaining 90 annotations (panel b), and variant 2 overlaps none of the ten upweighted annotations and all of the remaining 90 annotations (panel c). As the weighting constant increases, we observe that the weighted PINES p-value decreases for variant 1 and increases for variant 2. This depicts the tradeoff in power that can be used to select an appropriate weighting constant.

Regardless of whether a manual or enrichment-based weighting is employed to construct the matrix W, no annotation will be completely excluded from the model. For example in a study of pigmentation, the objective is for variants that have melanocyte-related annotations as well as exhibit evidence of functionality in other cell types to receive more significant scores than variants that only have melanocyte-related annotations. Another reason to rely on data from multiple cell types, even when the phenotypic effect of variants is limited to a single, well characterized cell type, is to gain statistical power from accumulating noisy correlated datasets.

### Comparison of PINES and annotation counting

To highlight the dynamics of weighted and unweighted PINES scores under different correlation structures of the annotation matrix, we simulate annotations under two different assumptions (Supplemental Fig. S14). In the first scenario, no correlation structure is present in the annotation space (panel a), so we generate 10,000 variants with 100 independent binary annotations (0/1). Under the second scenario, annotations exhibit correlation structure (panel d). Here we generate 10,0 variants with 10 annotation blocks, each containing 10 binary annotations (0/1). The annotations within each block are correlated with each other, but no correlation exists between annotations in different blocks. We highlight two variants: the variant that overlaps the first annotation from each of the 10 blocks of correlated annotations (green), and the variant that overlaps all ten annotations from the first block of correlated annotations (blue). Thus, both the green and the blue variant overlap ten annotations, but the selected annotations reflect different correlation patterns (uncorrelated annotations for green, correlated annotations for blue). We next designate the first annotation from the first correlation block as biologically relevant, and use this selection to explore the effect of weighting on PINES scores. In particular, both the green and the blue variant overlap the biologically relevant annotation. Panels c and f plot weighted PINES scores, with all variants that do not overlap the upweighted annotations are marked black.

A first point highlighted by Supplemental Fig. S14 is that, as opposed to raw counts, PINES scores are sensitive to correlation structure in the annotation space. Panel b shows almost equal PINES scores for both variants when no correlation structure is present in the annotation data. Under this scenario, PINES p-values and raw annotation counts will provide a similar ranking of the variants. Panel c shows that variants overlapping the upweighted annotation achieve more significant p-values than variants that do not overlap this annotation. When correlation structure is present, panel e shows the PINES and annotation counting deliver different results: the variant that overlaps a block of correlated annotations (blue) achieves less significant PINES p-values than the variants that overlaps multiple independent annotations (green). This distinction is not reflected by annotation counts, where both variants overlap a total of ten annotations. Panel f shows that weighted PINES scores also properly account for correlation structure. A second point highlighted by Supplemental Fig. S14 is that, as opposed to raw counts, PINES offers a principled method to perform annotations weighting. Panels c and f highlight the effect of annotation weights on PINES p-values under two different correlation structures of the annotation data.

### Other methods

A few approaches have been recently proposed to score noncoding regions and address the complexity of the annotation data in a principled manner. The Genome-Wide Annotation of Variants method (GWAVA) [10] aims to predict the impact of noncoding genetic variants based on a random forest classifier, using variants reported in the Human GeneMutation Database (HGMD) as deleterious training data, and common SNPs from the 1000 Genomes Project as benign examples. The CADD approach [11] is based on the premise that harmful mutations are edged out of the gene pool over time via natural selection and that variation that has not been selected against is thus less likely to be deleterious. Notable for CADD is that it uses a dataset of simulated mutations for training, which is then compared to observed variants. A score of deleteriousness is assigned to every possible SNP in the human genome. One of the most recent methods, Eigen-PC [9], is an unsupervised scoring framework that uses the eigen-decomposition of the covariance matrix associated with a collection of functional annotations to compute variant scores representing weighted sums of individual annotations. GenoCanyon [17] is a functional prediction method that performs unsupervised statistical learning using 22 computational and experimental annotations. Based on conservation scores and biochemical signals obtained from the ENCODE project a posterior probability of a genomic position being functional is computed. The authors mention cell type-specific scoring as a potential extension of GenoCanyon. DANN [12] is built upon the same feature set and training data as CADD, but instead of the linear kernel support vector machine used by CADD, DANN employs a deep neural network. An advantage of deep neural networks is their ability to capture non-linear relationships among features, and the authors show improved performance over CADD. LINSIGHT [14] is an unsupervised method that combines a generalized linear model for functional genomic data with a probabilistic model of molecular evolution to estimate the probability that individual mutations will have fitness consequences. FATHMM-MKL [13] integrates functional annotations from ENCODE with sequence conservation measures to predict the functional effects of genetic variants. FATHMM-MKL is a supervised method and relies on a positive training set of heritable germ-line mutations from the Human Gene Mutation Database (HGMD). DIVAN [16] is a supervised method that is trained disease by disease using annotated disease-specific variants. Due to the large number of epigenetic features but small number of training variants, feature selection is used to select the most informative set of features, and ensemble learning creates a more balanced risk/benign variant set in each base learner. In our comparisons of different methods we extracted pre-computed published scores for GWAVA, CADD, Eigen-PC, DANN, LINSIGHT, FATHMM-MKL, GenoCanyon, and DIVAN.

## Data access

PINES can be queried directly through a web interface at http://genetics.bwh.harvard.edu/pines/. The raw PINES source code is available on GitHub (https://github.com/PINES-scoring/PINES) and the annotation dataset is available on Zenodo (https://zenodo.org/record/1228512#.WuBft1MvyLI). The DOI for the annotation dataset is “10.5281/zenodo.1228512”. We include links to these resources on the PINES web interface, where users can directly compute weighted and unweighted PINES scores without additional computational overhead.

## Acknowledgments

We would like to thank Dr. Ivan Adzhubey for help with setting up the PINES web server.

